# Intercontinental prediction of soybean phenology via hybrid ensemble of knowledge-based and data-driven models

**DOI:** 10.1101/2020.09.22.306506

**Authors:** Ryan F. McCormick, Sandra K. Truong, Jose Rotundo, Adam P. Gaspar, Don Kyle, Fred van Eeuwijk, Carlos D. Messina

## Abstract

The timing of crop development has significant impacts on management decisions and subsequent yield formation. A large intercontinental dataset recording the timing of soybean developmental stages was used to establish ensembling approaches that leverage both discrete-time dynamical system models of soybean phenology and data-driven, machine-learned models to achieve accurate and interpretable predictions. We demonstrate that the knowledge-based, dynamical models can improve machine learning by generating expert-engineered features. Combining the predictions of the diverse component models via super learning resulted in a mean absolute error of 4.12 and 4.55 days to flowering (R1) and physiological maturity (R7), providing an improvement relative to the best benchmark model error of 6.90 and 15.47 days, respectively. The hybrid intercontinental model applies to a much wider range of management and temperature conditions than previous mechanistic models, enabling improved decision support as alternative cropping systems arise, farm sizes increase, and changes in the global climate continue to accelerate.

## Introduction

The timing of the changes in the life stages of a crop, also referred to as the crop’s phenology, represent primary determinants of the suitability of a crop for a growing region, its yield, and major management decisions such as planting date and agronomic treatments (Shaykewich, 1995). Allelic diversity in genetic pathways regulating crop phenology introduce significant variation in how, both within and between species, plants integrate environmental information and determine developmental stages. Models built to predict phenology for agronomic decision support are often relevant to only a limited set of prediction tasks such as a limited range of relative maturities, planting dates, locations, and/or photoperiod sensitivities, due to the inherent difficulty of building and training a broadly applicable and accurate model (dos Santos et al., 2019; Shaykewich and Bullock, 2018). However, as new sustainability practices and climate change motivate adoption of alternative management practices, such as planting in new target environments, seed increases in winter nurseries, or later planting of a shorter maturity crop in alternative cropping systems, more comprehensive approaches to phenology prediction are necessary.

Soybean is the world’s fourth largest crop as measured by area harvested and an important source of protein and oil (FAOSTAT 2016). As in most plants, the soybean life cycle is a well-regulated process that integrates environmental cues and internal states to determine the onset of phenological events. The timing of these events are the primary drivers of plant performance and reproductive success (Andrés and Coupland, 2012). Due to the economic and food security consequences that the timing of these events have, the understanding and prediction of soybean phenology has been a focus of research for decades (Brown, 1960; Cao et al., 2017; Hesketh et al., 1973; Shaykewich and Bullock, 2018; Wang et al., 1987). The present study builds off of this work by leveraging existing knowledge-based models of how the plant integrates key environmental determinants influencing phenology (Grimm et al., 1993; Jones et al., 2003; Salmerón and Purcell, 2016; Setiyono et al., 2007). Ultimately, this research sought to generate modeling strategies for predicting soybean phenology across disparate geographies and training procedures to generate an intercontinentally useful model.

Advances in computing and data science have driven massive increases in data availability and a concomitant increase in models describing biological systems, and models describing these systems vary in their purpose, accuracy, correctness, granularity, and interpretability. Prior to the increase in compute power, models of plant phenotypic outcomes given an environment typically existed either as parametric statistical models with explicit GxE interactions or as knowledge-based models composed of functions explicitly defining plant processes (Prusinkiewicz, 2004; Sinclair, 1986; van Eeuwijk et al., 2016). More recent efforts have focused on integrating GxE by coupling whole genome prediction with dynamical crop growth models, thereby generating phenotypic outcomes via non-linear functions of marker effects and environmental inputs (Cooper et al., 2016; Messina et al., 2018; Onogi et al., 2016; Technow et al., 2015). Purely data-driven approaches trained via machine learning have also garnered considerable attention as data generation continues to become more routine and higher throughput for plant systems (Liakos et al., 2018; Shakoor et al., 2019; Taghavi Namin et al., 2018). Parametric statistical models and process-based models can provide inferential and predictive ability in relatively data-poor environments and bring interpretability and understanding of the system being modeled, whereas in data-rich environments, data-driven machine-learned models often yield high predictive accuracy at the expense of interpretability. This tradeoff has motivated interest in hybrid modeling approaches in an effort to leverage the strengths of different modeling approaches and mitigate their respective weaknesses (Fan et al., 2015; Hamilton et al., 2017; Karpatne et al., 2017b, 2017a; Oyetunde et al., 2018; Pathak et al., 2018; Roberts et al., 2017).

While existing knowledge-based models can perform well under specific conditions or with laborious calibration, they are not sufficiently generalizable to support agronomic and breeding decisions across a variety of environmental conditions and continuous relative maturities (as opposed to discrete maturity groups). One possible approach to a general phenology model is the calibration or fitting of existing discrete-time dynamical systems models such as CROPGRO’s phenology module (hereafter referred to as CROPGRO for brevity) or SOYDEV (Boote et al., 1998; Jones et al., 2003; Salmerón and Purcell, 2016; Setiyono et al., 2007). However, re-fitting these models to new data often requires specialized experiments whose labor requirements restrict throughput to relatively few assays (Grimm et al., 1993). Like many biological models, these models are overparameterized, and they are unidentifiable given only the observations of calendar dates to developmental stages that are typically measured in applied field settings. This problem of fitting unidentifiable models with non-convex cost landscapes and limited data is not unique to the life sciences; while strategies exist to reparameterize the complex model to assist model fitting, crop growth modelers have generally employed various optimization approaches to regularize and fit the complex crop growth model (Archontoulis et al., 2014; He et al., 2010; Lamsal et al., 2017; Messina et al., 2018; Sexton et al., 2016; Wallach et al., 2001). The best optimization strategy will vary with the size and composition of the training dataset, sensitivity of the output to the selected set of input parameters, choice of priors and regularization schemes, and model runtime. Inspired by the success of machine learning approaches to identify optima (even if not the global optimum) that perform well in out-of-sample prediction tasks for models with large numbers of parameters (e.g., neural networks for image processing), this work develops a highly parallelized, gradient-free, multi-modal optimization strategy to explore high-dimensional parameter space and find satisfactory fits to available data (Kennedy, 2010; Spall, 1998; Whitley, 1994). This approach is used to recalibrate existing knowledge-based models given a large dataset of field observations, and the recalibrated models are shown to have improved prediction accuracy.

As an alternative to knowledge-based models, a second approach to a general phenology model is the utilization of advances in time-series modeling and model training made by the machine learning community. Specifically, artificial neurons that retain an internal state or memory can modulate their output based on past input; artificial neural networks built from such neurons can learn complex relationships between temporal sequences and output (e.g., speech to text applications). In other words, these networks have the potential to capture the sequence-dependent impact of environmental stimuli, such as a period of cool weather, on plant development. As such, network architectures including recurrent neural networks (RNN), like the Long Short Term Memory (LSTM) networks and their relatives, represent useful tools to model time-series data and predict phenological states (Hochreiter and Schmidhuber, 1997). These data-driven models have the potential to learn a mapping between daily inputs over time and the output phenological state on a given day, providing a data-driven analogue to the knowledge-based models mentioned previously. While the use of artificial neural networks to model soybean phenology is not new, the utility of a LSTM network to serve as a daily phenology model has not been explored to our knowledge (D. A. Elizondo et al., 1994; Zhang et al., 2009).

Moreover, it stands to reason that, since knowledge-based models represent a formal encapsulation of decades of research into system behavior, providing the prediction of a knowledge-based model as a feature should reduce the training burden on a machine-learned model and improve its predictive skill (Pathak et al., 2018). In this sense, knowledge-based models could be used to generate expert-engineered features from which a data-driven model could learn more effectively, and we explore this possibility in this paper.

The different model training approaches applied herein for the knowledge-based models and for the data-driven models do not have any guarantees regarding convergence on a global best fit, and their learning process is stochastic by nature. Therefore, instead of assuming that there exists a single “best” (i.e. most generalizable) model structure and parameters, model-selection approaches are eschewed for an ensembling technique that builds reliable meta-models given the predictions from the component models. This meta-model learns from the component model predictions, and is often referred to as a super learner. Most any regression procedure can serve as a super learner to stack the model predictions, and modern data science has successfully employed a variety of approaches from simple linear regression to artificial neural networks as super learners (Naimi and Balzer, 2018; Polley and van der Laan, 2010). In terms of providing uncertainty estimates regarding the prediction, Bayesian model averaging (BMA) and related approaches have emerged as practical ensembling techniques with useful properties in that it weighs individual models to provide some information on relative importance in the final ensemble prediction (Raftery et al., 2005; Yao et al., 2018).

In this work, an imbalanced, transcontinental soybean phenology dataset consisting of 13,673 records in 187 unique environments (defined here as unique combinations of planting date, latitude, and longitude) was used to demonstrate four concepts: (1) a multi-modal optimization strategy that can be used to train a crop growth model with a large dataset, (2) LSTM networks can be trained via machine learning to accurately predict soybean phenology, (3) knowledge-based model features can improve machine learning processes, and (4) combining the component model predictions via super learning reliably generates accurate out-of-sample predictions. The combined model applies to a wide variety of environments with an out-of-sample mean absolute error (MAE) of 4.12 and 4.55 days to R1 and R7, respectively, representing a 40% and 71% improvement over the best pre-existing model tested (6.90 and 15.47 days MAE) when averaged across the 10 folds. Accuracy for these and other key phenological stages is sufficient to direct management decisions and inform yield formation. We show that this approach yields considerable improvements in predictive power while still retaining some interpretability via the knowledge-based models to guide future experimentation and better understand system behavior underlying soybean phenology.

## Results

### (1) Dataset collection, evaluation of existing soybean phenology models, and model training overview

Data compilation from different public sources of relevant multi-environment trials and planting date experiments across years and a variety of latitudes was carried out and combined with data from trials internal to Corteva Agriscience. This resulted in 187 unique environments (defined as unique combinations of latitude, longitude, and planting date) with 13,673 records of observations of the calendar day of flowering (R1) and physiological maturity (R7) for a range of relative maturity groups (Fehr and Caviness, 1977). Of these 13,673 records, a subset also had observations for emergence and/or beginning seed filling (R5): 7,043 records had emergence, and 8,212 records had R5. 5,488 records included all four developmental stages. While additional phenological stages are characterized for soybean, the application for this study prioritized accurate predictions of R1 and R7, followed by R5 and emergence. The sampled environments range from −38.36° to 49.44° in latitude, covering a wide range of latitudes in growing regions of North and South America and representing approximately 83% of annual global soybean production; sampled lines range in relative maturity groups from −1.5 to 9.2 (i.e. less than MG 0 to greater than MG IX). Studies from which public data were obtained are listed in Supplemental Table 1.

A variety of soybean phenology models exist which generally serve to convert a time-series of daily temperatures and photoperiod to the calendar day of a developmental event, generally conditioned on a relative maturity group (Shaykewich and Bullock, 2018). To approximate the state of the art performance in soybean phenology modeling in the intercontinental dataset, the soybean phenology models described in CROPGRO and SOYDEV were re-implemented and applied without any calibration (Grimm et al., 1993; Jones et al., 2003; Salmerón and Purcell, 2016; Setiyono et al., 2007). After generalizing the discrete maturity parameters of the respective models to a continuous range of relative maturities (i.e. make a model parameter a function of continuous relative maturity instead of a discrete maturity group), both models demonstrated acceptable performance in some cases, but failed to generalize well across all environments and maturities (Figure 1). This is unsurprising, given that the range of photoperiods, temperatures, and maturities in the intercontinental dataset are outside of those sampled when the original model developers built their systems of equations and fit their model parameters. CROPGRO displayed useful performance for planting to R1 (MAE = 6.90 days), and exhibited substantial bias towards underprediction of days from planting to R7 (MAE = 15.47 days); SOYDEV displayed substantial error for both planting to R1 and planting to R7 (MAE = 10.59 and 25.14 days, respectively), particularly for maturities outside of those originally parameterized by Setiyono et al. (2007). As is generally the case, the models would need to be calibrated prior to use for decision support since predictions with their default parameters would be unsuitable for the range of environments and relative maturities present in the current prediction task.

**Figure 1.**
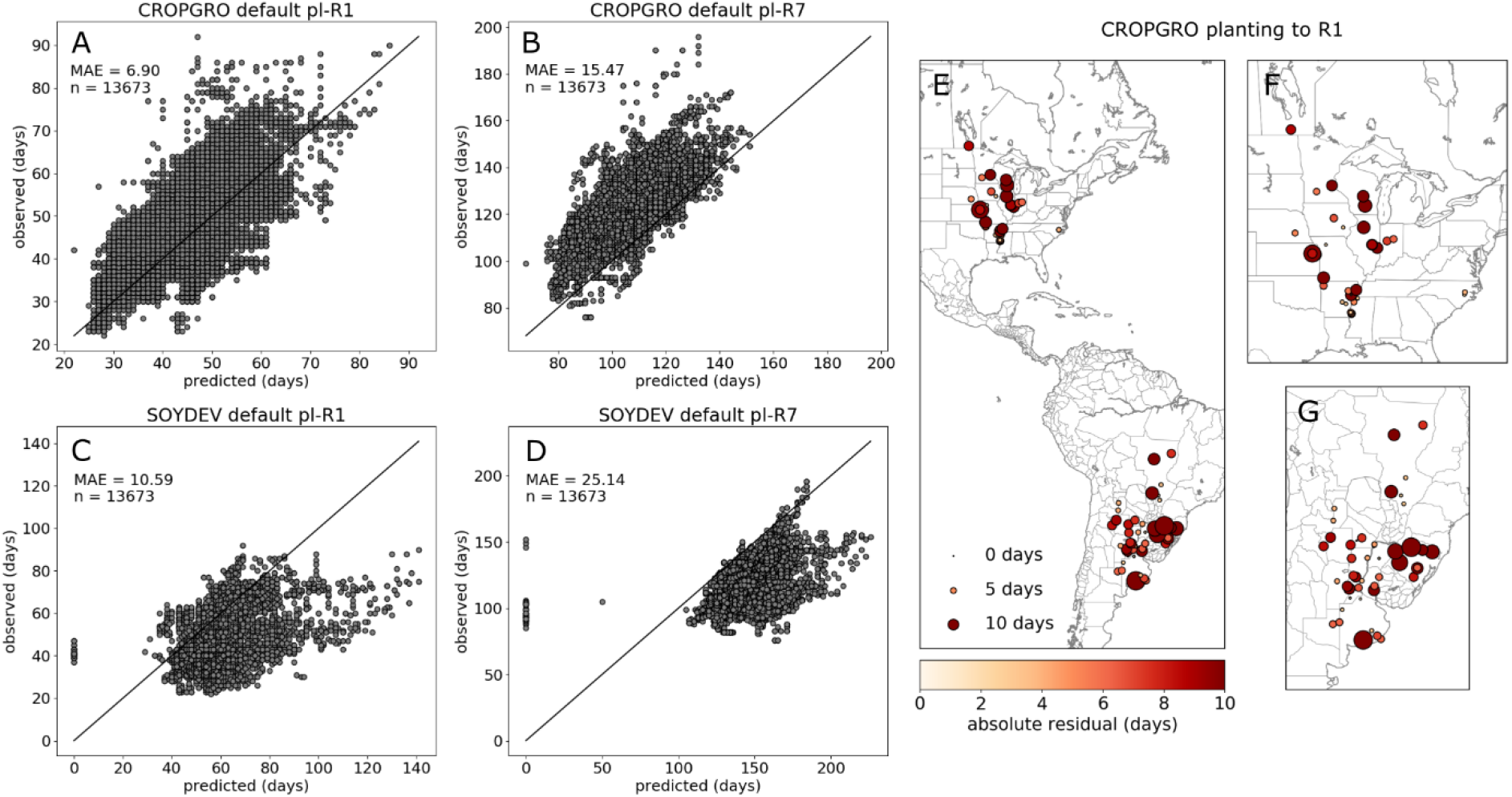
Performance of state of the art phenology models from CROPGRO and SOYDEV. Black line in scatter plots shows the one-to-one line for observed vs predicted for CROPGRO planting to R1 and planting to R7 (A, B) and likewise for SOYDEV (C, D). Predictions of zero by SOYDEV occurred due to relative maturities less than 0 failing to ever reach any stage, resulting in a prediction assigned as zero. For CROPGRO, default parameters generalized well for planting to R1 and R7 (A), though showed consistent underprediction for planting to R7 (B). Mean absolute residual by site using CROPGRO for planting to R1 (E, F, G). Residuals were not uniformly distributed with respect to factors like relative maturity, reflecting high model error when predicting cases outside of those the model was originally built for (data not shown).

Given the original dataset of 187 environments, ten independent, mutually exclusive Testing sets were generated on the basis of these environments. Each Testing set comprised 10% of the entire dataset, thereby creating 10 folds; the remaining 90% of environments in each fold were randomly divided into a Training and a Tuning set in a 60:40 split (Figure 2). The Training, Tuning, and Testing sets were generated on a per-environment basis, where a single environment could contain multiple records of phenological observations. Set partitioning was done on the basis of environments rather than records because, otherwise, models may have already learned using observations from the target environment; this represents the more trivial use case of predicting outcomes in an already observed environment. The Training and Tuning sets were used to fit component models and super learners, respectively; final prediction accuracy of the model training pipeline was evaluated on records from the Testing set environments.

**Figure 2.**
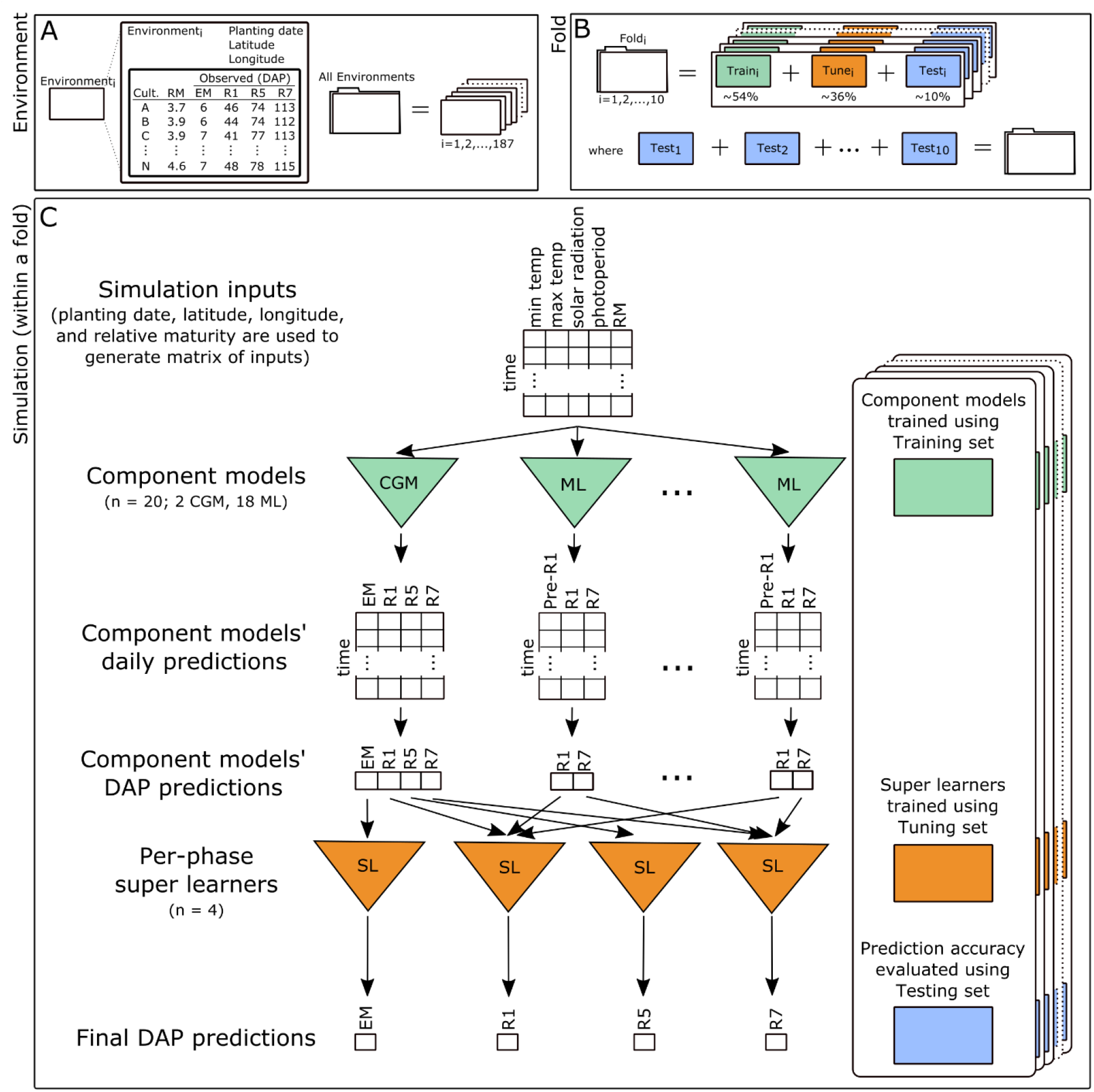
Graphical overview of data partitioning, training procedures, and processing pipeline. (A) Each environment comprises a unique latitude, longitude, and planting date associated with one or more records; each record corresponds to phenological observations or traits of days after planting (DAP) to reach a developmental stage for a soybean cultivar with a known relative maturity (RM). (B) 10 folds are generated by building 10 mutually exclusive Testing sets of environments, where each Testing set comprises 10% of the entire set of environments; the remaining 90% of the environments for each fold are randomly assigned to Training and Tuning sets on a 60:40 basis. (C) Overview of the processing pipeline for a single record. The record’s environment and its relative maturity define the matrix of daily inputs that component models receive. Component models, trained using environments from the Training set, generate daily outputs that are converted to a prediction of calendar days after planting (DAP) that a phenological phase is reached. Predictions made by component models are used as input to per-phase super learners that were trained using the Tuning set. The output of the super learners represents the final predictions; prediction accuracy was evaluated using the Testing set. Note that this does not depict the input of the crop growth models’ daily predictions to the machine-learned models, a process that is described below.

### (3) Training component models: Fitting knowledge-based phenology models

Fitting complex biological models to data remains a challenging problem, and a number of optimization strategies have been employed to obtain parameter estimates for crop growth models or their phenology models (Archontoulis et al., 2014; He et al., 2010; Lamsal et al., 2017; Messina et al., 2018; Sexton et al., 2016; Wallach et al., 2001). The approach used here assumes that there may be more than one equivalently good solution given the data (sometimes referred to as equifinality) and employs a highly-parallelized, multi-modal optimization strategy (Wong, 2015). In this manner, large populations of particles, each representing a parameter vector with a corresponding goodness of fit to the dataset, are used to explore the parameter space. Because the observations may fail to constrain the fit to a unique solution given the model structure, distinct particles may have similar goodness of fits. While this optimization strategy provides the opportunity to examine families of parameters that may cluster differently in parameter space but have equivalent fits, a single particle with the best fit (i.e. lowest cost) was chosen to represent the best model parameters given the Training set and used for subsequent analyses of a given fold. Thus, while the fit model generalizes well for prediction tasks, there are no guarantees that the variation explained by the best fitting model is driven by the biologically correct parameters, and the fits should either not be used for inference or used with caution. Most every parameter in the model, including parameters like optimal temperatures and photoperiod responses for different developmental phases, were jointly estimated, for a total of 32 free parameters in CROPGRO (Supplemental Methods). Of note, poor performance of both the default SOYDEV model and the re-fit SOYDEV model relative to CROPGRO led to subsequently dropping SOYDEV from further analyses.

Re-fitting the model resulted in improved out-of-sample performance across the 10 folds when predicting into the Tuning set, particularly for developmental stages past R1 where bias towards underprediction had been observed; the median across-fold MAE for the optimized model were 6.23 and 5.96 days to R1 and R7, respectively, and 6.59 and 14.69 days to R1 and R7 for the default model parameters (Figure 3). While the re-fit model has improved generalization error, it is notable that the prediction skill for R1 of the default model is performant, and that the prediction skill for R5 and R7 with the default model could be improved by a simple bias correction of the predictions; these suggest that the model structure and default parameters, while relatively simple, are capable of capturing large components of observed variation in the duration of developmental phases (Figure 1, Figure 3).

**Figure 3.**
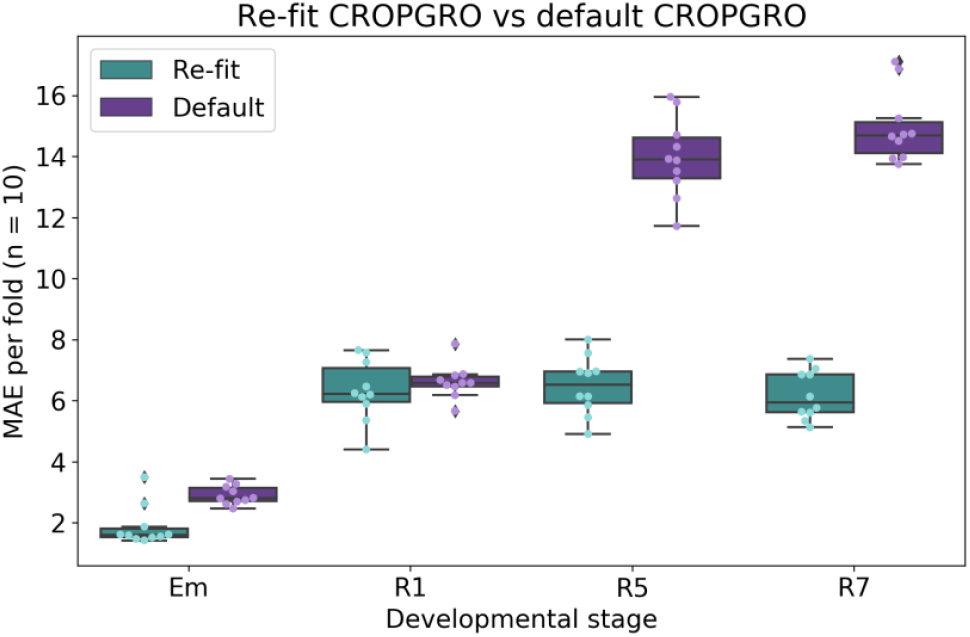
Comparison of out-of-sample accuracy for the re-fit CROPGRO with the default CROPGRO across 10 folds. In all cases, refitting the model using the Training set improved prediction accuracy in the out-of-sample Tuning set.

### (3) Training component models: Machine-learned models of soybean phenology using knowledge-based features

Recurrent neural network architectures such as Long Short Term Memory (LSTM) networks have demonstrated good performance in learning temporal or sequence dependent structure in data, and they could potentially serve as a data-driven analogue to the knowledge-based soybean phenology models. Thus, a training strategy was devised to train LSTM networks to predict daily soybean developmental stages and identify relevant hyperparameters for network architecture. LSTM networks were trained using inputs of minimum daily temperature, maximum daily temperature, solar radiation, night length as the complement of photoperiod, and relative maturity, where relative maturity was constant with respect to time (Figure 2). Thus, for each day in the sequence, the daily output of the LSTM network was a set of probabilities defining which of the discrete phenological phases the plant was likely in that day.

Hyperparameter tuning of neural networks is an outstanding challenge in machine learning, often accomplished by grid searches or evolutionary algorithms. A cursory grid search over the number of nodes and layers was performed using a Training set (and evaluating accuracy in a held-out subset of the Training set). A set of 9 different architectures (i.e. number of nodes and layers), including shallow and deep architectures, were retained. Once the architectures were defined, two separate models of each of these architectures with different random initializations were trained, generating a pool of 18 trained models of various architecture complexity within a fold. Notably, the relative performance for a given architecture varied such that there was no architecture that consistently outperformed any other; this observation is consistent with the premise of super learning that the combination of a collection of diverse models can generate more accurate predictions relative to using a single “best” model.

Since knowledge-based models represent a formal encapsulation of decades of research into the processes governing the system and are capable of explaining considerable variation in outcomes, their utility in machine learning was also examined. It stands to reason that providing this expert encapsulation of knowledge to a data-driven model as a feature should improve model training and generalization, so whether or not providing predictions from knowledge-based models as a feature to the data-driven model would improve performance was tested. To test this, the LSTM network architectures were trained using the aforementioned inputs only, or with the daily predictions from the default and re-fit CROPGRO model as additional features.

Inclusion of the predictions made by the CROPGRO models as features to the neural network model consistently improved the prediction accuracy of the collection of machine-learned models (Figure 4). The CROPGRO model outputs are simply functions of the same input data the neural network already received, and thus represent an expert-engineered feature. An interpretation of this result is that encapsulating prior knowledge as an engineered feature (i.e. the knowledge-based model’s prediction) improves the data-driven training process to fit more generalizable model (Figure 4). That is, despite the theoretical ability of neural networks to faithfully represent arbitrary mapping functions, an imperfect training process identifies a poor local optimum. Expert features in the form of the knowledge-based model’s prediction assists the training process to better find improved, more generalizable models. The median MAE for models trained without CROPGRO features were 4.84 and 5.92 days for planting to R1 and planting to R7, respectively, whereas the median MAE for models trained with CROPGRO features were 4.69 and 5.54 days for planting to R1 and planting to R7, respectively. Subsequent analyses were performed using the data-driven models that used the knowledge-based models’ predictions as inputs.

**Figure 4.**
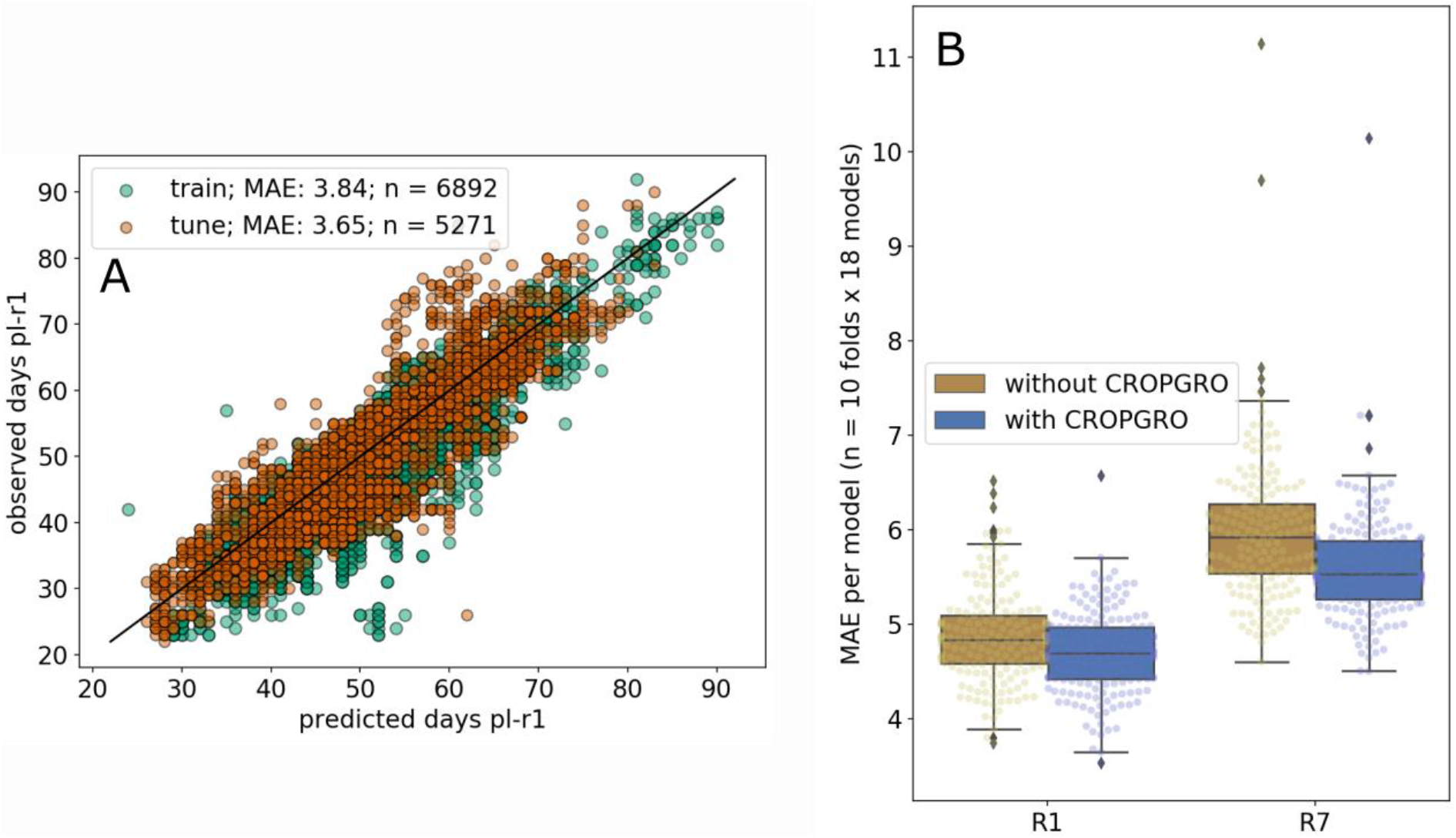
Data-driven approaches can learn temporal dynamics to predict phenology, and they are improved by predictions from knowledge-based models. (A) Example of a well-performing model for planting to R1 of one initialization of one LSTM network architecture from one fold (where the out-of-sample Tuning set MAE corresponds to one “R1 with CROPGRO” point in panel B). (B) Comparison of out-of-sample performance in the Tuning set for the collection of machine-learned models trained using the base environmental features alone (labeled “without CROPGRO”) or with both the default CROPGRO and optimized CROPGRO predictions as additional features (labeled “with CROPGRO”). The median performance of the entire collection of models is improved by inclusion of the knowledge-based model predictions during training even though the knowledge-based model is simply a transform of the same environmental inputs already provided to the data-driven model.

Unlike the knowledge-based models, the LSTM network models were trained only to predict R1 and R7. This was performed in order to maximize the amount of data the networks could be trained on, as there were considerably fewer observations containing all stages, and the handling of missing data in the context of training a recurrent neural network is not well established. As such, the neural networks were not trained to predict emergence or R5; the knowledge-based models generate predictions for those stages in subsequent analyses.

### (4) Combining component model predictions with super learning

Given a pool of models, model selection techniques are often used to choose a single “best” model for prediction and inference. Alternatively, since the best model for inference may not be the best model for prediction, model ensembling approaches such as meta-regression or super learning have gained popularity for prediction tasks; these approaches takes advantage of model diversity to improve predictions (Breiman, 1996; Naimi and Balzer, 2018). The simplest ensembling approach is an averaging of the predictions of every model in the pool, such that each individual model’s prediction is equally weighted in the final ensemble prediction. For applications where the relative utility of individual models is of interest, Bayesian model averaging (BMA) has emerged as a useful approach capable of weighing models, thereby providing some indication of which models are performing best while also leveraging their combined information (Hoeting et al., 1999; Raftery et al., 2005).

The pool of component models consisted of 18 data-driven and 2 knowledge-based models, all of which operated on daily time steps (Figure 2). For each model and for each developmental stage that the model predicted, the number of calendar days between planting and the developmental stage (DAP) was obtained. All models predicted R1 and R7, whereas only the knowledge-based models predicted emergence and R5. In a conceptually similar approach to Raftery et al., (2005), the records from the Tuning set were used to regress the observations on the predictions to obtain a bias corrected Bayesian linear regression model, generating a pool of Bayesian linear regression models that were then ensembled and weighted via stacking of predictive distributions described by Yao et al. (2018).

For predictions of R1 and R7, the super learner had access to the full pool of 20 models, whereas emergence and R5 only contained the default CROPGRO and re-fit CROPGRO models in their pool (Figure 2). Median out-of-sample MAE for predictions into the Testing set across the 10 folds were compared for (i) the super learner, (ii) the simple ensemble point average of each model’s predictions (i.e., all models weighted equally), (iii) CROPGRO with optimized parameters, and (iv) CROPGRO with default parameters (Figure 5). For each stage, the median across-fold MAE of the final predictions from the super learners was 1.37, 4.12, 5.55, and 4.55 days for planting to emergence, to R1, to R5, and to R7, respectively.

**Figure 5.**
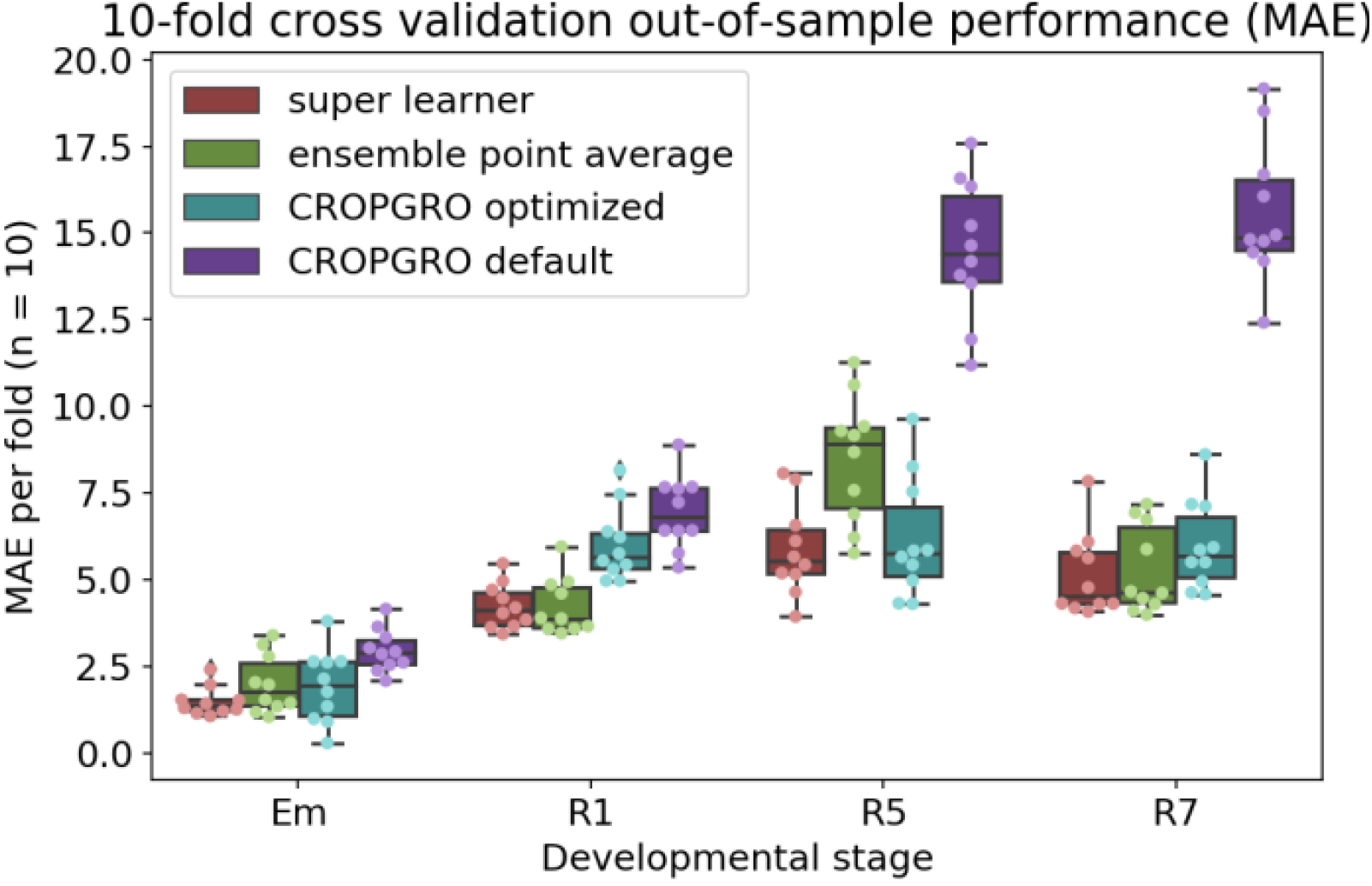
Comparison of out-of-sample performance across folds for the super learners, the ensemble point average (i.e., all models weighted equally), CROPGRO with optimized parameters, and CROPGRO with default parameters. Ensemble models for emergence (Em) and R5 only contain the optimized and default CROPGRO, whereas ensemble models for R1 and R7 additionally contain the 18 neural network models. In this case, out-of-sample performance is evaluated on only the Testing set of each fold.

As expected, all approaches that had been trained using a subset of the dataset had improved prediction accuracy for the out-of-sample Testing set relative to CROPGRO with default parameters. In all cases, the super learners performed at parity or better than the ensemble point averages (i.e. all models’ point prediction weighted equally), with the added benefit of identifying which models could be reasonably excluded from the ensemble. Because many of the models in the pool are given no weight by the super learner, this procedure has the significant benefit of retaining the predictive accuracy of the whole ensemble but reducing the number of models that need to be maintained and their deployment footprint for production settings. Similarly, in model deployment and production settings, this also facilitates the examination of potential tradeoffs between accurate models with larger memory footprints and runtimes vs those with smaller footprints and runtimes (e.g., for deployment on a mobile device). Ultimately, the super learners resulted in a median out-of-sample MAE of 4.12 and 4.55 days to R1 and R7, respectively, providing sufficient accuracy for decision support in a number of applications.

## Discussion

Fundamentally, all phenotypes are a consequence of genotype by environment interactions by nature of the genome’s integration of environmental signals. Historically, breeding progress has been driven by efforts that model genetic effects as largely independent of the environment, and this approach is particularly successful when the target population of environments displays relatively little environmental variation. However, as data generation and computational capacity continues to advance, extensions of these traditional approaches have been made that formalize the knowledge that overt phenotypes are non-linear transforms of genetic states and the environment over time (Messina et al., 2018). This work further builds on these premises, seeking to maximize the ability to transfer information learned about how given genetics interact with environments in one geography (e.g., late maturities in Brazil) to another (e.g., the United States).

Data acquisition in the life sciences will continue to present challenging modeling tasks for the foreseeable future, where *p* >> *n* due to the complexity of the system and relative inability to experimentally sample it (where *p* is the number of model parameters and *n* is the number of observations). Some measurements can be readily obtained at low costs (e.g., marker genotypes), whereas others are more expensive and laborious (e.g., multi-environment trials, longitudinal data), generating both data-rich and data-poor settings from which actionable conclusions need to be drawn. Moreover, even in data-rich environments where machine-learned models can perform well, there is a drive to bring improved interpretability to these models or to constrain them with prior knowledge. (Hazard et al., 2019; Marcus, 2018). The current work hybridizes the modeling strategies by training data-driven models with using knowledge-based predictions as features, as well as ensembling the knowledge-based and data-driven predictions by training a meta-model; considerable work remains to create truly hybrid systems that are easily used, capable of providing system understanding and inference, and identify novel components of the system (Fan et al., 2015; Hamilton et al., 2017, 2015; Karpatne et al., 2017b). These interpretable and adaptive models will enable both understanding and enable prescriptive decision support, and may continue to develop in the form of probabilistic programming and model-based machine learning (Bishop, 2013; Salvatier et al., 2016). Moreover, as super learners become more sophisticated, it stands to reason that they will be able to learn to use the signal processing provided by individual component models as abstractions to build hierarchical representations of the world, not unlike the human mind (Iten et al., 2018; Marcus, 2018).

Given that the neural network models outperform the optimized CROPGRO, it suggests that our current understanding of the system may need to be adapted to incorporate aspects of the system that the data-driven approach captures that the knowledge-based model does not. That said, that the performance improvement of the super learner over the default model is not more drastic is a testament to the body of knowledge developed regarding soybean phenology over the past decades. This body of knowledge helped to define the relevant features used as input to the data-driven model, namely temperature and photoperiod. Additionally, it is notable that the addition of CROPGRO predictions as features for the data-driven approach enabled the training procedure to generally identify better optima than the environmental features alone across an entire collection of tested network architectures. This implies that the knowledge-based model provided an informative integration of the environmental features that the training procedure for the LSTM was otherwise unable to identify.

This work builds on a long history of research into the genetic basis of and modeling of soybean phenology, and represents a practical milestone in integrating some of this information. However, the current work assumes that the genetic variation in soybean phenology can be compressed into a single scalar: the relative maturity. While the model achieves acceptable accuracy for the target applications, additional work should examine the utility of additional genetic information, such as markers for the soybean E-loci or markers for stem growth habit to distinguish between determinate and indeterminate lines (Messina et al., 2006; Tian et al., 2010). Future work will also benefit from advances in remote sensing efforts to accurately measure phenological events (Zeng et al., 2016), as well as advances in optimization that enable efficient searches over parameter spaces, hyperparameter spaces, and optimization strategies for data-driven models (Li and Malik, 2016).

The end application of the model is to support decisions across a wide variety of geographies by enabling growers and agronomists a forecast to plan crop management events, including planting and harvest dates, pesticide applications, and irrigation events, as well as to examine how maturities planted outside of their typical zone will perform. As the world’s geopolitics and climate continue to change, accurate prediction of outcomes for agricultural systems outside of the norm will be critical for rapid adaptation.

## Methods

### (1) Soybean phenology and weather database

The soybean phenology database used in this study was derived from public and private sources of data, with phenology stages based on Fehr and Caviness (1977). Phenology data for North America was derived predominantly from Corteva Agriscience’s internal resources. Data from Argentina was obtained from publicly available soybean tests coordinated by the National Institute of Agricultural Technology (INTA). Data on soybean phenology from Brazil were derived from available technical reports. A list of the public sources of observations included in the analysis is available in Supplemental Table 1. A proprietary Corteva weather repository and interpolation system was used to acquire daily historical weather observations from public and private weather stations for North and South America at the experimental coordinates.

### (2) Data harmonization and partitioning

The data were partitioned on the basis of unique environments, defined as unique latitude, longitude, and planting date (defined by month, day, and year) combinations to ensure that trained models were not trained using records that shared the same environment with out-of-sample records; here, a record is defined as group of phenological observations (i.e. traits) for a relative maturity in an environment. It could be argued that different planting dates within a site-year have sufficiently correlated weather to cause information leakage between in-sample and out-of-sample records and lead to over-optimistic out-of-sample accuracy metrics, but we consider the photoperiods to be sufficiently different to merit consideration as unique environments. Some data sources recorded observations of full maturity (R8) instead of physiological maturity (R7); while factors like relative humidity, temperature, wind speed, and rainfall can introduce variation in the duration between R7 and R8 across different environments, the duration can be practicably treated as a constant number of calendar days for this application (Gaspar et al., 2017; Martinez-Feria et al., 2017). Thus, observations of R8 were harmonized to R7 prior to any analyses by subtracting a constant 9 calendar days, and the harmonized observations were considered equivalent to R7 observations. The dataset consisted of 187 unique environments with a total of 13,673 records. All 13,673 records contained planting date, flowering (R1) date, and physiological maturity (R7) date. A subset of 7,043 records also contained emergence date, 8,212 recorded beginning seed filling (R5) date, and 5,488 records included all four developmental stages.

The 187 environments were partitioned in to Training, Tuning, and Testing sets in 10 folds (Figure 2). Four environments were permanently assigned to the Training set due to representing outlier combinations of environment and relative maturities. For the remaining 183 environments, each fold withheld a random selection of 10% of the environments as the Testing set, where the environments in each fold’s Testing set were mutually exclusive of all other folds. The Testing set of the first 9 folds contained 18 environments, and the final 10^th^ fold contained 21. The remaining environments in a fold were then randomly split on a 60:40 basis into a Training and Tuning set, with 103 environments in the Training set and 66 in the Tuning set (fold 10 contained 63 in the Tuning set).

Individual component models in the model pool or library (i.e., the collection of models whose predictions were provided to the super learners) were trained using records from the Training set. The super learners were trained using the component models’ predictions and records from the Tuning set. The performance of the trained super learners was evaluated using the Testing set (Figure 2).

### (3) Knowledge-based phenology model implementation and optimization

The phenology models for soybean from CROPGRO and SOYDEV were re-implemented with OpenCL and interfaced with PyOpenCL to enable highly parallelized execution of the models (Jones et al., 2003; Klöckner et al., 2012; Setiyono et al., 2007; Stone et al., 2010). A swarm of particles, each representing a parameter vector, was used to explore the parameter space, where each particle moved down an approximated gradient using simultaneous perturbation stochastic approximation (SPSA) in parallel to find a satisfactory minimum of the cost function (Spall, 1998). In brief, SPSA takes a random subset of the parameters and generates two new parameter vectors, one resulting from a small positive adjustment to the subset of parameters and one resulting from a small negative adjustment. The cost of both are evaluated, and the parameters are updated in the direction that minimizes cost. For this application, the cost was defined as the negative log-likelihood of the parameter vector given the data, where missing observations (e.g., missing R5 date) did not contribute to cost. In this manner, all observations for a record (e.g., planting to emergence, R1, R5, and R7) could be used simultaneously to fit the dynamical model.

A given parameter vector (i.e. a particle) would be updated according to SPSA using randomized minibatches over the records; once all particles finished all minibatches, the worst (i.e. highest cost or worst fit) particles were repopulated with parameters that were recombinations of the best particles’ parameters. The space searched for each parameter was bounded by a defined range (Supplemental Methods). The search was initialized by gridding a swarm of 4,800 particles across the parameter space, and performing parameter updates on minibatches of size 500. The bounds of the range a given parameter could occupy were manually assigned, and per-parameter learning rates were automatically assigned using the magnitude of the range to be searched for the parameter. Forty rounds of evolution were performed, meaning that each of the 4,800 particles completed all minibatches and had the potential to be recombined 40 times. If a training set had 7,500 records, this would lead to 4,800 * (7,500 * 2) * 40 simulations, or 2.88 billion simulations explored during training. While this multi-modal optimization strategy allows the exploration of equifinality and similarly behaving families of parameters, the parameter vector with the minimum cost found at the end of any round of evolution was retained and used as the set of optimized parameters for subsequent analyses.

### (4) Machine-learned models and super learners

All machine learned models were varying architectures of LSTM networks trained using Keras and Tensorflow (Abadi et al., 2016; Chollet and others, 2015). Models were trained using daily inputs of minimum temperature, maximum temperature, night length, solar radiation, relative maturity group (which was constant with respect to time), and an indicator variable to indicate whether planting had occurred or not to predict the developmental stage reached that day (as an integer). Model architectures ranged from shallow architectures with one hidden layer and 64 nodes, to deeper architectures with 2 or 3 hidden layers and up to 2,024 nodes using a tanh activation function; the output layer used a softmax activation function. Categorical cross entropy was used as the loss function, models were trained using Adam as the optimization strategy for up to 50 epochs with early stopping, and the best performing model on the Tuning set out of the epochs was retained for each architecture, generating a population of 18 models for each fold.

Bayesian model averaging as implemented in PyMC3 was used to integrate the predictions of the component models (Raftery et al., 2005; Salvatier et al., 2016; Yao et al., 2018). First, for each component model, a bias-corrected linear model for each developmental stage was fit using the number of calendar days from planting to the developmental stage derived from the model’s predictions as input and the observed calendar days as output using the Tuning set to estimate the posterior (Raftery et al., 2005). For emergence and R5 this involved only the default and optimized CROPGRO models; for R1 and R7 this involved those two models and the additional 18 trained neural network architectures. Stacking of predictive distributions given the population of these bias-corrected models was performed for each developmental stage to find an optimal weighing of component models for each stage (Yao et al., 2018). That is, different stages have different subsets of models contributing to the final predictions. Final performance was evaluated on the Testing set (Figure 2).

## Acknowledgements

The authors wish to acknowledge Marie Dolleman, Scott Birkett, Kirk Hansen, Riley McDowell, Duke Takle for their technical contributions in application hosting and productionization.

## Supplemental Material

**Supplemental Table 1.** Public data sources from which soybean phenological data were obtained.

| **Public source** | **Country** | **Observations (n)** |
| --- | --- | --- |
| Gaspar et al. 2017. Crop Sci. 57:2170–2182 | USA | 60 |
| Fundação MS - Resultados de Experimentação e Campos Demonstrativos de Soja - Safra 2014/2015 | Brazil | 204 |
| Fundação MS - Resultados de Experimentação e Campos Demonstrativos de Soja - Safra 2015/2016 | Brazil | 474 |
| Avaliação do desempenho produtivo de diferentes cultivares de soja em Nova Mutum – MT, na safra 2016 / 2017 | Brazil | 95 |
| Fundação Pro-sementes - Desempenho de cultivares de soja indicadas para Rio Grande do Sul - 2017/2018 | Brazil | 372 |
| Ensaio de competição de cultivares de soja 2014 to 2016 – MT, Estação Rural Técnica | Brazil | 257 |
| Red Nacional de Evaluación de Cultivares de Soja (RECSO) - 2014/2015 | Argentina | 2910 |
| Red Nacional de Evaluación de Cultivares de Soja (RECSO) - 2015/2016 | Argentina | 1769 |
| Red Nacional de Evaluación de Cultivares de Soja (RECSO) - 2016/2017 | Argentina | 1156 |

## Supplemental Methods

### Phenology model and re-fitting

The CROPGRO phenology model was implemented in OpenCL for parallelized optimization. Briefly, each developmental phase (emergence to R1, R1 to R5, R5 to R7) requires a target number of cumulative photothermal units to advance to the next phase. Under optimal conditions (i.e. optimal temperature and photoperiod), the number of photothermal units accumulated on a day would be one; otherwise it is a fraction of a unit.

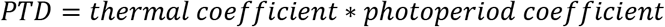

Where the PTD, fraction of a photothermal day, is the product of the thermal coefficient and photoperiod coefficient, graphically shown below. Thus, under optimal development conditions, the number of cumulative PTD to advance phases is equal to the number of calendar days; otherwise, the number of calendar days needed is greater than the PTD threshold.

**Supplemental Methods Figure 1.**
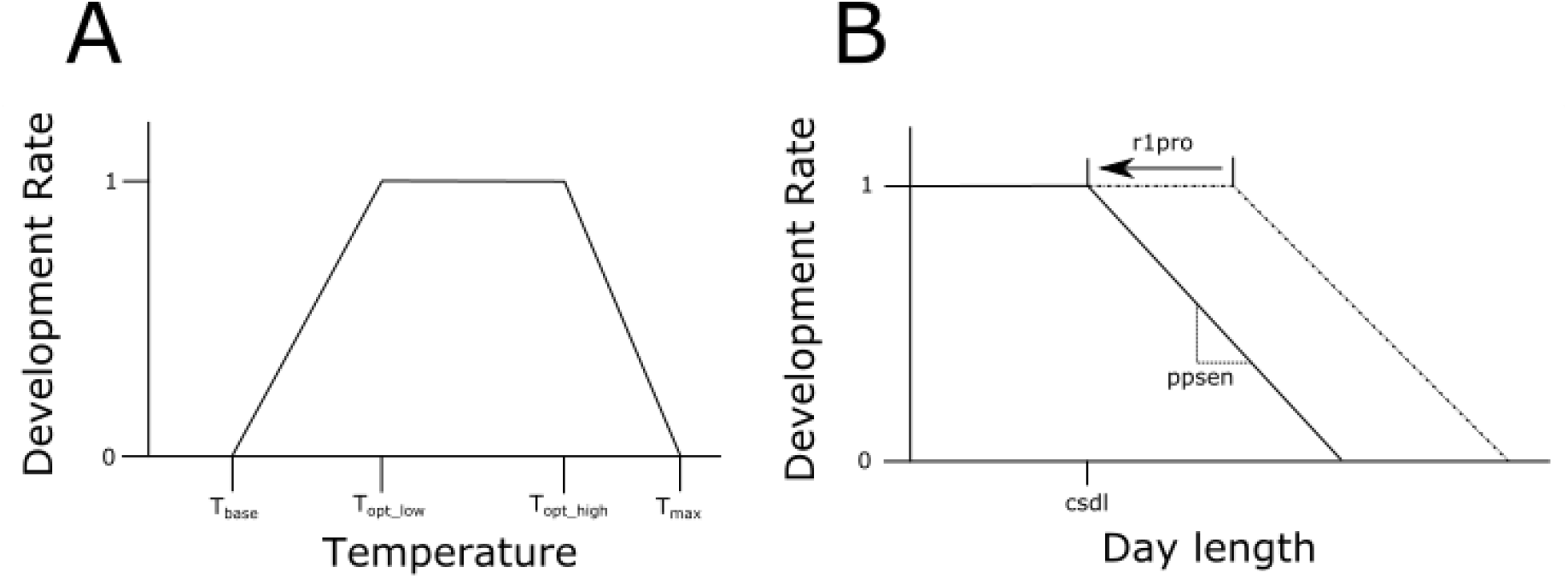
Thermal (A) and photoperiod (B) functions used to determine how many PTD would be accumulated on a given day. During optimization, it is possible that some parameters take on values that render other parameters irrelevant. For example, if the maximum temperature (T_max_) takes on a value less than the optimum high temperature (T_opt_high_), then T_opt_high_ becomes irrelevant, as the development rate becomes zero above T_max_.

For each developmental stage, the number of cumulative PTD required to advance to the next stage is a function of maturity group. Based on the relationship between the default values provided by CROPGRO and maturity group, the cumulative PTD threshold was modeled as a polynomial to obtain PTD as a function of relative maturity. Emergence to R1 was a first order, R1 to R5 was a fourth order, and R5 to R7 was a second order polynomial. For refitting parameters, the intercept (i.e. number of PTD for relative maturity 0 to advance its phase under optimal conditions) was assigned as the minimum number of calendar days observed in the dataset for a relative maturity 0 observation to advance through the phase. As such, 7 free parameters were estimated to determine cumulative PTD thresholds for each phase.

For each developmental stage, the thermal coefficient, calculated as piecewise linear function based on a minimum base temperature, an optimal temperature lower bound, an optimal temperature upper bound, and a maximum base temperature (graphically depicted above). As such, 16 free parameters (T_base_, T_opt_low_, T_opt_high_, and T_max_ each for each phase: planting to emergence, emergence to R1, R1 to R5, and R5 to R7) were fit for temperature responses.

The photoperiod response dictated by the critical short day length (csdl) and photoperiod sensitivity (ppsen) were both modeled as third order polynomials with respect to maturity group, and constant across developmental phases. The reduction in csdl after anthesis (r1pro) was treated as constant across maturity groups and developmental phases, leading to a total of 9 free parameters. Of note, planting to emergence was not impacted by the photoperiod response function.

The cost of a parameter vector was calculated as the negative log probability of the parameters given the available observations for a record assuming a normal distribution, and this cost was minimized.

**Supplementary Methods Table 1.** Table of default parameters, a set of re-fit parameters corresponding to one fold’s best fit, and the parameter boundaries within which the initialized parameter vectors were gridded and not permitted to explore beyond. Note that, due to the fitting strategy employed, fitted parameters should not be considered reliable for inference; e.g., note that some of the table functions have a maximum base temperature that render the maximum optimum temperature irrelevant.

| Parameter indexing string | Default | Re-Fit | Lower bound explored | Upper bound explored |
| --- | --- | --- | --- | --- |
| R1PRO | 0.279 | -1.458581 | -2 | 2 |
| CSDL_FIRSTORDER_COEF | -0.24227 | -0.397733 | -1 | 1 |
| CSDL_SECONDORDER_COEF | 0 | 0.014179 | -0.1 | 0.1 |
| CSDL_THIRDORDER_COEF | 0 | 0.000377 | -0.01 | 0.01 |
| CSDL_INTERCEPT | 14.07409 | 14.83173 | 12 | 16 |
| PPSEN_FIRSTORDER_COEF | 0.053835 | 0.02108 | -1 | 1 |
| PPSEN_SECONDORDER_COEF | -0.00683 | 0.001629 | -0.1 | 0.1 |
| PPSEN_THIRDORDER_COEF | 0.000329 | 0.000157 | -0.001 | 0.001 |
| PPSEN_INTERCEPT | 0.166189 | 0.009315 | 0 | 1 |
| PLEM_INTERCEPT | 3.6 | 3.6 | Fixed | Fixed |
| PLEM_MINBASETEMP | 7 | 0.627509 | -40 | 20 |
| PLEM_MAXBASETEMP | 45 | 78.21355 | 30 | 80 |
| PLEM_MINOPTTEMP | 28 | 53.88043 | 15 | 90 |
| PLEM_MAXOPTTEMP | 35 | 25.22636 | 20 | 80 |
| EMR1_FIRSTORDER_COEF | 0.678182 | 0.898344 | -10 | 10 |
| EMR1_INTERCEPT | 16.42727 | 16.42727 | Fixed | Fixed |
| EMR1_MINBASETEMP | 7 | 6.635806 | -40 | 20 |
| EMR1_MAXBASETEMP | 45 | 79.04411 | 30 | 80 |
| EMR1_MINOPTTEMP | 28 | 32.35435 | 15 | 90 |
| EMR1_MAXOPTTEMP | 35 | 47.47038 | 20 | 80 |
| R1R5_FIRSTORDER_COEF | -0.44503 | 0.634236 | -1 | 1 |
| R1R5_SECONDORDER_COEF | 0.449301 | -0.174032 | -1 | 1 |
| R1R5_THIRDORDER_COEF | -0.06663 | -0.021729 | -0.1 | 0.1 |
| R1R5_FOURTHORDER_COEF | 0.002914 | 0.003754 | -0.01 | 0.01 |
| R1R5_INTERCEPT | 13.02098 | 13.02098 | Fixed | Fixed |
| R1R5_MINBASETEMP | 6 | -20.23749 | -40 | 20 |
| R1R5_MAXBASETEMP | 45 | 76.97128 | 30 | 80 |
| R1R5_MINOPTTEMP | 26 | 61.50523 | 15 | 90 |
| R1R5_MAXOPTTEMP | 30 | 78.94601 | 20 | 80 |
| R5R7_FIRSTORDER_COEF | 1.001259 | 0.310288 | -2 | 2 |
| R5R7_SECONDORDER_COEF | -0.02867 | -0.006204 | -1 | 1 |
| R5R7_INTERCEPT | 30.61539 | 30.61539 | Fixed | Fixed |
| R5R7_MINBASETEMP | -15 | -17.79777 | -40 | 20 |
| R5R7_MAXBASETEMP | 45 | 39.977 | 30 | 80 |
| R5R7_MINOPTTEMP | 26 | 36.31358 | 15 | 90 |
| R5R7_MAXOPTTEMP | 34 | 74.33067 | 20 | 80 |
| **cost** | 527544.8 | 184794.5 |  |  |
| **MAE_target_0** | 3.17488 | 1.976063 |  |  |
| **MAE_target_1** | 7.316628 | 6.050265 |  |  |
| **MAE_target_2** | 14.36696 | 7.212268 |  |  |
| **MAE_target_3** | 16.33564 | 6.481427 |  |  |
| **numRecords** | 6784 | 6784 |  |  |

## Literature cited

Abadi, M., Barham, P., Chen, J., Chen, Z., Davis, A., Dean, J., Devin, M., Ghemawat, S., Irving, G., Isard, M., Kudlur, M., Levenberg, J., Monga, R., Moore, S., Murray, D.G., Steiner, B., Tucker, P., Vasudevan, V., Warden, P., Wicke, M., Yu, Y., Zheng, X., 2016. TensorFlow: A system for large-scale machine learning, in: 12th USENIX Symposium on Operating Systems Design and Implementation (OSDI 16). pp. 265–283.

Andrés, F., Coupland, G., 2012. The genetic basis of flowering responses to seasonal cues. Nat. Rev. Genet. 13, 627–639. https://doi.org/10.1038/nrg3291

Archontoulis, S.V., Miguez, F.E., Moore, K.J., 2014. A methodology and an optimization tool to calibrate phenology of short-day species included in the APSIM PLANT model: application to soybean. Environ. Model. Softw. 62, 465–477. https://doi.org/10.1016/j.envsoft.2014.04.009

Bishop, C.M., 2013. Model-based machine learning. Philos. Trans. R. Soc. Math. Phys. Eng. Sci. 371, 20120222. https://doi.org/10.1098/rsta.2012.0222

Boote, K.J., Jones, J.W., Hoogenboom, G., Pickering, N.B., 1998. The CROPGRO model for grain legumes, in: Tsuji, G.Y., Hoogenboom, Gerrit, Thornton, P.K. (Eds.), Understanding Options for Agricultural Production, Systems Approaches for Sustainable Agricultural Development. Springer Netherlands, Dordrecht, pp. 99–128. https://doi.org/10.1007/978-94-017-3624-4_6

Breiman, L., 1996. Stacked regressions. Mach. Learn. 24, 49–64. https://doi.org/10.1007/BF00117832

Brown, D.M., 1960. Soybean Ecology. I. Development-temperature relationships from controlled environment studies. Agron. J. 52, 493–496. https://doi.org/10.2134/agronj1960.00021962005200090001x

Cao, D., Takeshima, R., Zhao, C., Liu, B., Jun, A., Kong, F., 2017. Molecular mechanisms of flowering under long days and stem growth habit in soybean. J. Exp. Bot. 68, 1873–1884. https://doi.org/10.1093/jxb/erw394

Chollet, F., others, 2015. Keras.

Cooper, M., Technow, F., Messina, C., Gho, C., Totir, L.R., 2016. Use of crop growth models with whole-genome prediction: application to a maize multienvironment trial. Crop Sci. 56, 2141–2156. https://doi.org/10.2135/cropsci2015.08.0512

D. A. Elizondo, R. W. McClendon, G. Hoogenboom, 1994. Neural network models for predicting flowering and physiological maturity of soybean. Trans. ASAE 37, 981–988. https://doi.org/10.13031/2013.28168

dos Santos, C., Salmerón, M., Purcell, L.C., 2019. Soybean phenology prediction tool for the US midsouth. Agric. Environ. Lett. 4. https://doi.org/10.2134/ael2019.09.0036

Fan, X.-R., Kang, M.-Z., Heuvelink, E., de Reffye, P., Hu, B.-G., 2015. A knowledge-and-data-driven modeling approach for simulating plant growth: A case study on tomato growth. Ecol. Model. 312, 363–373. https://doi.org/10.1016/j.ecolmodel.2015.06.006

Fehr, W., Caviness, C., 1977. Stages of soybean development. Spec. Rep.

Gaspar, A.P., Laboski, C.A.M., Naeve, S.L., Conley, S.P., 2017. Dry matter and nitrogen uptake, partitioning, and removal across a wide range of soybean seed yield levels. Crop Sci. 57, 2170–2182. https://doi.org/10.2135/cropsci2016.05.0322

Grimm, S.S., Jones, J.W., Boote, K.J., Hesketh, J.D., 1993. Parameter estimation for predicting flowering date of soybean cultivars. Crop Sci. 33, 137–144. https://doi.org/10.2135/cropsci1993.0011183X003300010025x

Hamilton, F., Berry, T., Sauer, T., 2015. Predicting chaotic time series with a partial model. Phys. Rev. E 92, 010902. https://doi.org/10.1103/PhysRevE.92.010902

Hamilton, F., Lloyd, A.L., Flores, K.B., 2017. Hybrid modeling and prediction of dynamical systems. PLOS Comput. Biol. 13, e1005655. https://doi.org/10.1371/journal.pcbi.1005655

Hazard, C.J., Fusting, C., Resnick, M., Auerbach, M., Meehan, M., Korobov, V., 2019. Natively interpretable machine learning and artificial intelligence: preliminary results and future directions. arXiv 1901.00246v2.

He, J., Jones, J.W., Graham, W.D., Dukes, M.D., 2010. Influence of likelihood function choice for estimating crop model parameters using the generalized likelihood uncertainty estimation method. Agric. Syst. 103, 256–264. https://doi.org/10.1016/j.agsy.2010.01.006

Hesketh, J.D., Myhre, D.L., Willey, C.R., 1973. Temperature control of time intervals between vegetative and reproductive events in soybeans. Crop Sci. 13, 250–254. https://doi.org/10.2135/cropsci1973.0011183X001300020030x

Hochreiter, S., Schmidhuber, J., 1997. Long short-term memory. Neural Comput. 9, 1735–1780. https://doi.org/10.1162/neco.1997.9.8.1735

Hoeting, J.A., Madigan, D., Raftery, A.E., Volinsky, C.T., 1999. Bayesian model averaging: a tutorial. Stat. Sci. 14, 382–401.

Iten, R., Metger, T., Wilming, H., del Rio, L., Renner, R., 2018. Discovering physical concepts with neural networks. arXiv arXiv:1807.10300v2.

Jones, J.W., Hoogenboom, G., Porter, C.H., Boote, K.J., Batchelor, W.D., Hunt, L.A., Wilkens, P.W., Singh, U., Gijsman, A.J., Ritchie, J.T., 2003. The DSSAT cropping system model. Eur. J. Agron., Modelling Cropping Systems: Science, Software and Applications 18, 235–265. https://doi.org/10.1016/S1161-0301(02)00107-7

Karpatne, A., Atluri, G., Faghmous, J.H., Steinbach, M., Banerjee, A., Ganguly, A., Shekhar, S., Samatova, N., Kumar, V., 2017a. Theory-guided data science: a new paradigm for scientific discovery from data. IEEE Trans. Knowl. Data Eng. 29, 2318–2331. https://doi.org/10.1109/TKDE.2017.2720168

Karpatne, A., Watkins, W., Read, J., Kumar, V., 2017b. Physics-guided neural networks (PGNN): an application in lake temperature modeling.

Kennedy, J., 2010. Particle swarm optimization, in: Sammut, C., Webb, G.I. (Eds.), Encyclopedia of Machine Learning. Springer US, Boston, MA, pp. 760–766. https://doi.org/10.1007/978-0-387-30164-8_630

Klöckner, A., Pinto, N., Lee, Y., Catanzaro, B., Ivanov, P., Fasih, A., 2012. PyCUDA and PyOpenCL: A scripting-based approach to GPU run-time code generation. Parallel Comput. 38, 157–174. https://doi.org/10.1016/j.parco.2011.09.001

Lamsal, A., Welch, S.M., Jones, J.W., Boote, K.J., Asebedo, A., Crain, J., Wang, X., Boyer, W., Giri, A., Frink, E., Xu, X., Gundy, G., Ou, J., Arachchige, P.G., 2017. Efficient crop model parameter estimation and site characterization using large breeding trial data sets. Agric. Syst. 157, 170–184. https://doi.org/10.1016/j.agsy.2017.07.016

Li, K., Malik, J., 2016. Learning to optimize. arXiv arXiv:1606.01885v1.

Liakos, K.G., Busato, P., Moshou, D., Pearson, S., Bochtis, D., 2018. Machine learning in agriculture: a review. Sensors 18, 2674. https://doi.org/10.3390/s18082674

Marcus, G., 2018. Deep learning: a critical appraisal.

Martinez-Feria, R., Archontoulis, S.V., Licht, M.A., 2017. How fast do soybeans dry down in the field? [WWW Document]. URL https://crops.extension.iastate.edu/cropnews/2017/09/how-fast-do-soybeans-dry-down-field (accessed 1.14.20).

Messina, C.D., Jones, J.W., Boote, K.J., Vallejos, C.E., 2006. A gene-based model to simulate soybean development and yield responses to environment. Crop Sci. 46, 456–466. https://doi.org/10.2135/cropsci2005.04-0372

Messina, C.D., Technow, F., Tang, T., Totir, R., Gho, C., Cooper, M., 2018. Leveraging biological insight and environmental variation to improve phenotypic prediction: Integrating crop growth models (CGM) with whole genome prediction (WGP). Eur. J. Agron. 100, 151–162. https://doi.org/10.1016/j.eja.2018.01.007

Naimi, A.I., Balzer, L.B., 2018. Stacked generalization: an introduction to super learning. Eur. J. Epidemiol. 33, 459–464. https://doi.org/10.1007/s10654-018-0390-z

Onogi, A., Watanabe, M., Mochizuki, T., Hayashi, T., Nakagawa, H., Hasegawa, T., Iwata, H., 2016. Toward integration of genomic selection with crop modelling: the development of an integrated approach to predicting rice heading dates. Theor. Appl. Genet. 129, 805–817. https://doi.org/10.1007/s00122-016-2667-5

Oyetunde, T., Bao, F.S., Chen, J.-W., Martin, H.G., Tang, Y.J., 2018. Leveraging knowledge engineering and machine learning for microbial bio-manufacturing. Biotechnol. Adv. 36, 1308–1315. https://doi.org/10.1016/j.biotechadv.2018.04.008

Pathak, J., Wikner, A., Fussell, R., Chandra, S., Hunt, B.R., Girvan, M., Ott, E., 2018. Hybrid forecasting of chaotic processes: using machine learning in conjunction with a knowledge-based model. Chaos Interdiscip. J. Nonlinear Sci. 28, 041101. https://doi.org/10.1063/1.5028373

Polley, E., van der Laan, M., 2010. Super learner in prediction. UC Berkeley Div. Biostat. Work. Pap. Ser.

Prusinkiewicz, P., 2004. Modeling plant growth and development. Curr. Opin. Plant Biol. 7, 79–83. https://doi.org/10.1016/j.pbi.2003.11.007

Raftery, A.E., Gneiting, T., Balabdaoui, F., Polakowski, M., 2005. Using bayesian model averaging to calibrate forecast ensembles. Mon. Weather Rev. 133, 1155–1174. https://doi.org/10.1175/MWR2906.1

Roberts, M.J., Braun, N.O., Sinclair, T.R., Lobell, D.B., Schlenker, W., 2017. Comparing and combining process-based crop models and statistical models with some implications for climate change. Environ. Res. Lett. 12, 095010. https://doi.org/10.1088/1748-9326/aa7f33

Salmerón, M., Purcell, L.C., 2016. Simplifying the prediction of phenology with the DSSAT-CROPGRO-soybean model based on relative maturity group and determinacy. Agric. Syst. 148, 178–187. https://doi.org/10.1016/j.agsy.2016.07.016

Salvatier, J., Wiecki, T.V., Fonnesbeck, C., 2016. Probabilistic programming in Python using PyMC3. PeerJ Comput. Sci. 2, e55. https://doi.org/10.7717/peerj-cs.55

Setiyono, T.D., Weiss, A., Specht, J., Bastidas, A.M., Cassman, K.G., Dobermann, A., 2007. Understanding and modeling the effect of temperature and daylength on soybean phenology under high-yield conditions. Field Crops Res. 100, 257–271. https://doi.org/10.1016/j.fcr.2006.07.011

Sexton, J., Everingham, Y., Inman-Bamber, G., 2016. A theoretical and real world evaluation of two Bayesian techniques for the calibration of variety parameters in a sugarcane crop model. Environ. Model. Softw. 83, 126–142. https://doi.org/10.1016/j.envsoft.2016.05.014

Shakoor, N., Northrup, D., Murray, S., Mockler, T.C., 2019. Big data driven agriculture: big data analytics in plant breeding, genomics, and the use of remote sensing technologies to advance crop productivity. Plant Phenome J. 2. https://doi.org/10.2135/tppj2018.12.0009

Shaykewich, C.F., 1995. An appraisal of cereal crop phenology modelling. Can. J. Plant Sci. 75, 329–341. https://doi.org/10.4141/cjps95-057

Shaykewich, C.F., Bullock, P.R., 2018. Modeling soybean phenology. Agroclimatol. Link. Agric. Clim. agronomymonogra. https://doi.org/10.2134/agronmonogr60.2018.0002

Sinclair, T.R., 1986. Water and nitrogen limitations in soybean grain production I. Model development. Field Crops Res. 15, 125–141. https://doi.org/10.1016/0378-4290(86)90082-1

Spall, J.C., 1998. Implementation of the simultaneous perturbation algorithm for stochastic optimization. IEEE Trans. Aerosp. Electron. Syst. 34, 817–823. https://doi.org/10.1109/7.705889

Stone, J.E., Gohara, D., Shi, G., 2010. OpenCL: a parallel programming standard for heterogeneous computing systems. Comput. Sci. Eng. 12, 66–73. https://doi.org/10.1109/MCSE.2010.69

Taghavi Namin, S., Esmaeilzadeh, M., Najafi, M., Brown, T.B., Borevitz, J.O., 2018. Deep phenotyping: deep learning for temporal phenotype/genotype classification. Plant Methods 14, 66. https://doi.org/10.1186/s13007-018-0333-4

Technow, F., Messina, C.D., Totir, L.R., Cooper, M., 2015. Integrating crop growth models with whole genome prediction through approximate bayesian computation. PLOS ONE 10, e0130855. https://doi.org/10.1371/journal.pone.0130855

Tian, Z., Wang, X., Lee, R., Li, Y., Specht, J.E., Nelson, R.L., McClean, P.E., Qiu, L., Ma, J., 2010. Artificial selection for determinate growth habit in soybean. Proc. Natl. Acad. Sci. 107, 8563–8568. https://doi.org/10.1073/pnas.1000088107

van Eeuwijk, F.A., Bustos-Korts, D.V., Malosetti, M., 2016. What should students in plant breeding know about the statistical aspects of genotype × environment interactions? Crop Sci. 56, 2119–2140. https://doi.org/10.2135/cropsci2015.06.0375

Wallach, D., Goffinet, B., Bergez, J.-E., Debaeke, P., Leenhardt, D., Aubertot, J.-N., 2001. Parameter estimation for crop models. Agron. J. 93, 757–766. https://doi.org/10.2134/agronj2001.934757x

Wang, J., McBlain, B.A., Hesketh, J.D., Woolley, J.T., Bernard, R.L., 1987. A data base for predicting soybean phenology. Biotronics 16, 25–38.

Whitley, D., 1994. A genetic algorithm tutorial. Stat. Comput. 4, 65–85. https://doi.org/10.1007/BF00175354

Wong, K.-C., 2015. Evolutionary multimodal optimization: a short survey. arXiv arXiv:1508.00457v1.

Yao, Y., Vehtari, A., Simpson, D., Gelman, A., 2018. Using stacking to average Bayesian predictive distributions (with discussion). Bayesian Anal. 13, 917–1007. https://doi.org/10.1214/17-BA1091

Zeng, L., Wardlow, B.D., Wang, R., Shan, J., Tadesse, T., Hayes, M.J., Li, D., 2016. A hybrid approach for detecting corn and soybean phenology with time-series MODIS data. Remote Sens. Environ. 181, 237–250. https://doi.org/10.1016/j.rse.2016.03.039

Zhang, J.-Q., Zhang, L.-X., Zhang, M.-H., Watson, C., 2009. Prediction of soybean crowth and development using artificial neural network and statistical models. Acta Agron. Sin. 35, 341–347. https://doi.org/10.1016/S1875-2780(08)60064-4

